# Low genomic divergence and high gene flow between locally adapted populations of the swamp sparrow

**DOI:** 10.1101/359992

**Authors:** P. Deane-Coe, R. Greenberg, I. J. Lovette, R. G. Harrison

## Abstract

Populations that have recently diverged across sharp environmental gradients provide an opportunity to study the mechanisms by which natural selection drives adaptive divergence. Inland and coastal populations of the North American swamp sparrow have become an emerging model system for studies of natural selection because they are morphologically and behaviourally distinct despite a very recent divergence time (<15,000 years), yet common garden experiments have demonstrated a genetic basis for their phenotypic differences. We characterized genomic patterns of variation within and between inland and coastal swamp sparrows via reduced representation sequencing in order to reconstruct the contributions of demography, gene flow and selection to this case of recent adaptive divergence. Compared to inland swamp sparrows, coastal swamp sparrows exhibited fewer polymorphic sites and reduced nucleotide diversity at those sites, indicating that a bottleneck and/or recent selective sweeps occurred in that population during coastal colonization and local adaptation. Estimates of genome-wide differentiation (F_ST_=0.02) and sequence divergence (**Φ**_ST_=0.05) between inland and coastal populations were very low, consistent with postglacial divergence. A small number of SNPs were strongly differentiated (max F_ST_=0.8) suggesting selection at linked sites. Swamp sparrows sampled from breeding sites at the habitat transition between freshwater and brackish marshes exhibited high levels of genetic admixture. Such evidence of active contemporary gene flow makes the evolution and maintenance of local adaptation in these two populations even more notable. We summarize several features of the swamp sparrow system that may facilitate the maintenance of adaptive diversity despite gene flow, including the presence of a magic trait.

## Introduction

Birds offer many striking examples of local adaptation, particularly in terms of two evolutionarily labile traits: bill shape and plumage. Both traits are capable of substantial phenotypic change in response to natural selection, over both shallow and deep phylogenetic timescales (eg. Kusmierski et al. 1997; Lovette et al. 2002; Burns et al. 2002). In many cases, bill or plumage traits likely evolve via post-speciation selection, when allopatric taxa that are already reproductively isolated experience different forms of selection on these phenotypes depending on their environment (Dobzhansky 1940). Alternatively, when natural selection itself is responsible for driving genetic divergence between lineages during the evolution of local adaptation, we refer to the process as adaptive divergence. Since adaptive divergence may lead to the evolution of reproductive isolation (“ecological speciation”; Mayr 1963; Harrison 1991; Rice and Hostert 1993; Funk 1998; Schluter 2000, 2001; Coyne and Orr 2004; Nosil et al. 2005; Rundle and Nosil 2005; Nosil 2012), case studies of the early stages of adaptive divergence offer an opportunity to understand the mechanisms by which natural selection drives diversification over time. How does selection create and maintain locally adapted lineages before reproductive isolation has evolved between them?

Ecologically distinct populations of the North American swamp sparrow (*Melospiza georgiana*) have served as a natural laboratory for studies of adaptive divergence for decades. In the eastern USA there is both a widespread inland form of the swamp sparrow (*M. g. georgiana*), and a range-restricted coastal form (*M. g. nigrescens*) that breeds in brackish tidal marshes of the Delaware and Chesapeake Bays (Figure 1; Greenberg and Droege 1990; Beadell et al. 2003). “Coastal Plain” swamp sparrows were first recognized as distinct from inland swamp sparrows more than 60 years ago (Bond and Stewart 1951), and various phenotypes that distinguish the two forms have since been studied extensively. The coastal population differs from inland populations in at least ten traits (Table 1), including a deeper bill and melanic plumage (Figure 2; Greenberg and Droege 1990). Field studies have demonstrated that at least eight of the ten derived traits that distinguish coastal swamp sparrows are adaptive in coastal environments (Table 1; Goldstein et al. 2004; Grenier and Greenberg 2005; Olsen et al. 2008, 2013; Peele et al. 2009; Greenberg et al. 2012), and common garden rearing experiments have confirmed that there is a strong genetic basis to most trait differences between the populations, including bill and plumage (Table 1; Ballentine and Greenberg 2010). Natural selection for deeper bills in coastal habitats has, in turn, biomechanically constrained the range of possible bandwidths at which coastal males can trill (Ballentine 2006), driving further population divergence through nonrandom mating. Inland swamp sparrow females prefer to mate with males that sing broad-bandwidth songs and maximize vocal performance, but coastal swamp sparrow females prefer the songs of coastal males, who compensate for a loss of bandwidth by increasing trill rate (Ballentine et al. 2013a, 2013b). Swamp sparrows are the third empirical example of bill shape as a “magic trait” (Gavrilets 2004): natural selection on the bills of Darwin’s finches and crossbills also imposes biomechanical constraints on song, facilitating divergence in this mating signal and driving the evolution of reproductive isolation (Podos 2001; Podos et al. 2004; Huber and Podos 2006; Huber et al. 2007; Smith and Benkman 2007).

**Table 1.** Proposed adaptive hypotheses for eight traits that distinguish coastal Swamp Sparrows from inland forms, and their empirical support from field studies. Trait differences indicated by an asterisk (*) persist when coastal and inland birds are reared in a common garden. Clutch size could not be measured in captivity (Ballentine and Greenberg 2010).

| Trait | Hypotheses | Proposed selective agent | Proposed adaptive benefit | Support |
| --- | --- | --- | --- | --- |
| ↑ Bill depth* | *Trophic adaptation* | More abundant invertebrate prey in coastal habitat | Improves prey capture<sup>1,2,3</sup> | some |
|  | *Abiotic adaptation* | Heat stress due to exposed, water-limited habitats | Convective radiative heat loss<sup>4</sup> | Y |
| ↑ Bill dimorphism | *Thermal budgets* | Territorial defense causes heat stress in males | Convective radiative heat loss<sup>4</sup> | some |
|  | *Intraspecific divergence* | High breeding densities, competition for resources | Sex-based diet specialization<sup>5</sup> | N |
|  | *Intrasexual selection* | High breeding densities, territorial competition | Competitive advantage<sup>5</sup> | unclear |
|  | *Female choice* | High breeding densities, competition for paternity | Females prefer larger bills<sup>3</sup> | Y |
| ↑ Plumage melanism* | *Predation adaptation* | Anoxia and iron sulphides cause darker sediments | Crypsis<sup>1</sup> | untested |
|  | *Feather preservation* | Feather degrading bacteria thrive in saline habitats | Confers resistance to bacteria<sup>6</sup> | Y |
|  | *Intrasexual selection* | High breeding densities, territorial competition | Larger black plumage badges<sup>7</sup> | Y |
| ↓ Plumage rustiness* | *Predation adaptation* | Anoxia and iron sulphides cause darker sediments | Crypsis<sup>1</sup> | untested |
|  | *Evolutionary constraint* | Selection for plumage melanism | None, females prefer rusty crowns<sup>7,8</sup> | Y |
| Delayed molt, extended breeding* | *Predation adaptation* | High predation on coast causes nest failure | Allows re-nesting<sup>9</sup> | some |
|  | *Abiotic adaptation* | Coastal habitats experience late spring flooding | Facilitates delayed breeding<sup>1</sup> | some |
| ↓ Clutch size | *Predation adaptation* | Nest predation rates are higher in coastal habitats | Reduces provisioning, detectability<sup>9</sup> | Y |
|  | *Longevity trade-off* | Lifetime breeding effort favored over current | Improves annual survivorship<sup>9</sup> | N |
|  | *Abiotic adaptation* | Temperature extremes, egg loss during laying | Shorter laying avoids extreme events<sup>9</sup> | Y |
| ↓ Song bandwidth* | *Evolutionary constraint* | Larger bills constrain the bandwidth of male trills | None, broad bandwidth favored<sup>10,11,12</sup> | Y |
| ↑ Song trill rate | *Acoustic adaptation* | Faster trills transmit better in open habitats | Improves signal transmission<sup>13,14</sup> | some |
|  | *Female choice* | Faster songs are honest indicators of quality | Females prefer fast trills<sup>10,12,15,16</sup> | Y |
|  | *Signal compensation* | Bill size constrains bandwidth, males trill faster | Females prefer challenging songs<sup>10,11</sup> | some |

1. Greenberg and Droege 1990 2. Grenier and Greenberg 2005 3. Olsen et al. 2013 4. Greenberg et al. 2012 5. Greenberg and Olsen 2010 6. Peele et al. 2009 7. Olsen et al. 2008a 8. Olsen et al. 2010 9. Olsen et al. 2008b 10. Ballentine 2006 11. Ballentine et al. 2013a 12. Liu et al. 2008 13. Morton 1975 14. Wiley and Richards 1978 15. Ballentine 2009 16. Ballentine et al. 2013b

**Figure 1.**
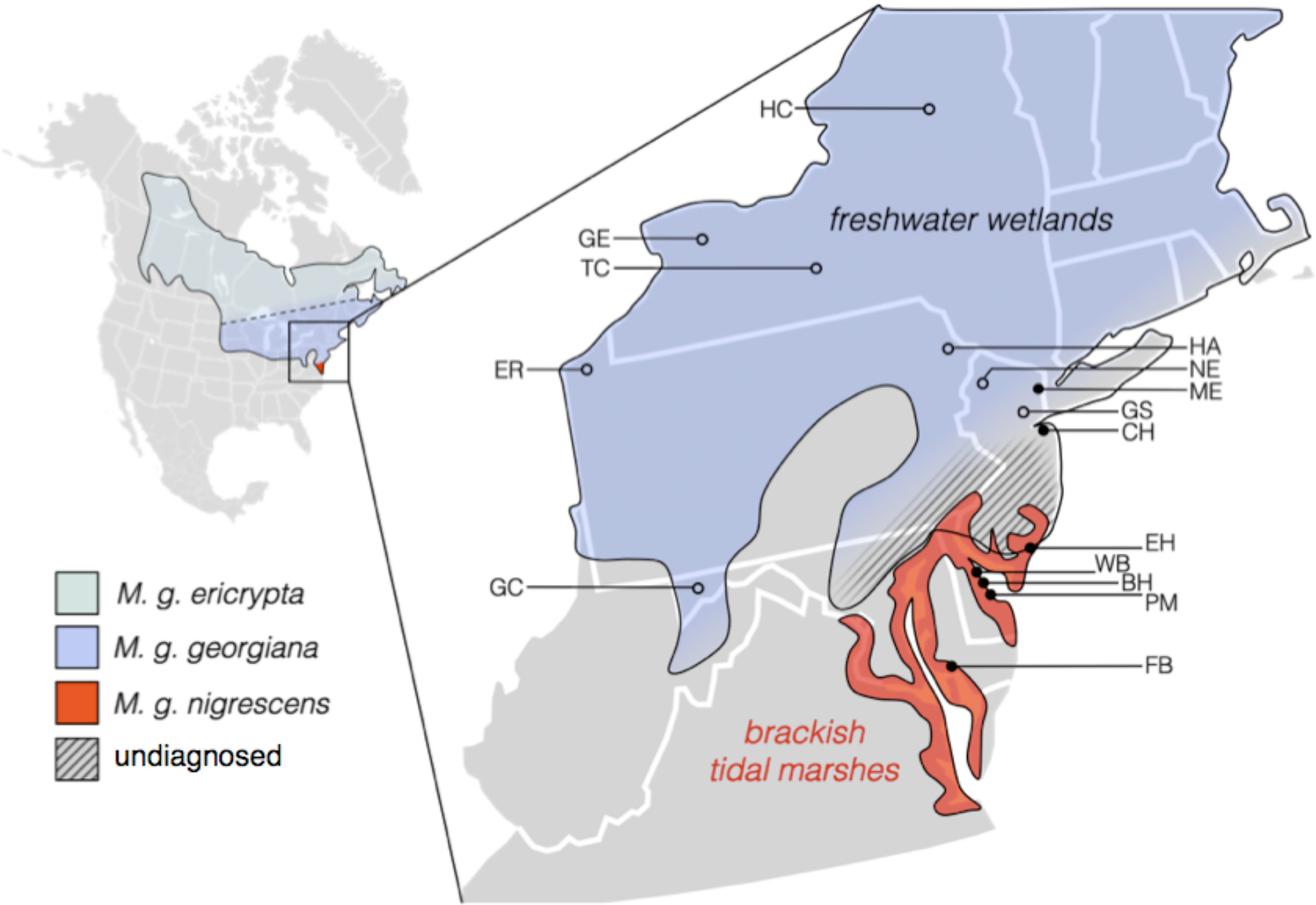
Breeding range of Swamp Sparrow subspecies (Greenberg and Droege 1990; Beadell et al. 2003). Freshwater (°) and brackish (•) sampling sites are shown with two-letter site codes (Table 2). Hatching denotes an area of very low breeding density in which the phenotypes of birds have not been studied. Dashed line denotes the approximate southern extent of the boreal subspecies (*ericrypta*), not included in this study.

**Figure 2.**
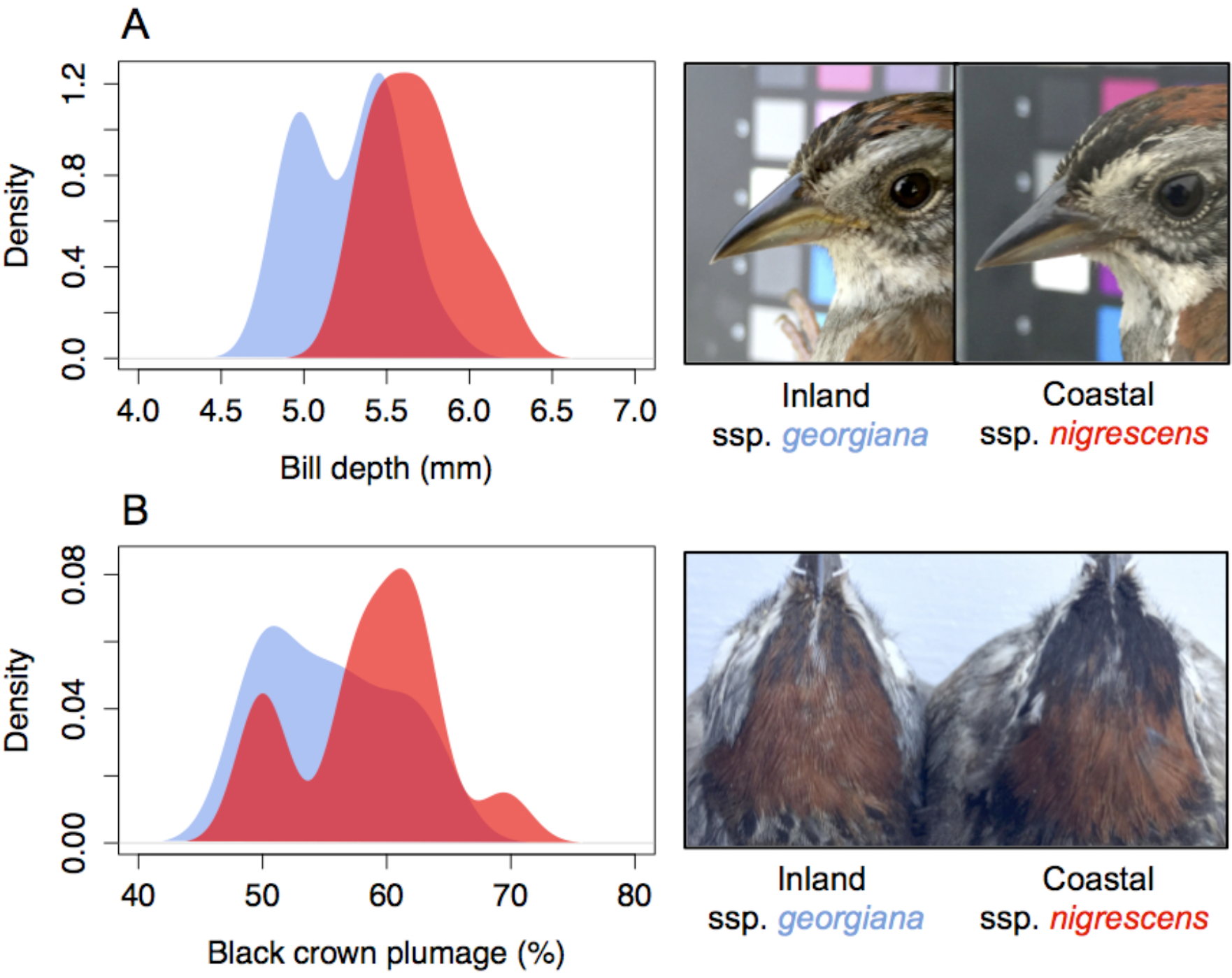
Bill depth (A) and the extent of black crown plumage (B) are two of many traits that subtly distinguish inland (blue) and coastal (red) swamp sparrows. Morphological data include only birds from allopatric sites (N=29) and are from Greenberg and Droege (1990). Photos are of breeding males (inland: Hamilton County NY and Assunpink NJ, coastal: Fishing Bay MD).

Although many of the adaptive traits in coastal swamp sparrows have a heritable genetic basis, coastal birds have not had much time to evolve them. Coastal tidal marshes did not exist in the northeastern or mid-atlantic regions during the Last Glacial Maximum (LGM) because northeastern coasts were under the ice sheet, and mid-atlantic coasts were hydrologically unstable due to glacial outflow from rivers and streams (Malamud-Roam et al. 2006). Tidal marsh habitats also require sediment accretion from rising sea levels in order to establish (Pethick 1984; Warren and Niering 1993). Coastal swamp sparrows have therefore likely colonized and adapted to this novel habitat rapidly, within the last 15,000 years. Consistent with a recent divergence, inland and coastal swamp sparrows have proven indistinguishable using allozymes and mitochondrial DNA (Balaban 1988; Greenberg et al. 1998). A recent study using microsatellite markers was the first to successfully resolve subtle genetic differentiation between coastal and inland swamp sparrows, and a genetic contact zone in northern New Jersey coincident with the ecotone between brackish and freshwater marshes (Greenberg et al. 2016).

To reconstruct the evolutionary processes responsible for adaptive divergence between inland and coastal swamp sparrows, we quantify genome-wide molecular variation for the species and extract patterns of historical demography, gene flow and selection. We also test for an association between phenotypic and genomic divergence, and characterize levels of admixture in the putative inland/coastal hybrid zone with genome-level sampling.

## Methods

### Sample collection and phenotypic measurements

We compiled 92 DNA samples from male and female swamp sparrows from 15 sites distributed across the breeding range of inland and coastal forms in the northeastern US, including samples from the putative contact zone in northern New Jersey (Figure 1; Tables 2, S1). Most were blood samples collected from birds banded in the 2001 breeding season (N=66), and the remainder were blood samples from banded birds or tissue samples from vouchered specimens, collected by P. Deane-Coe (P.D.) or other collectors from the Cornell University Museum of Vertebrates during the 2007-2014 breeding seasons (N=26). We collected four voucher specimens and six blood samples from a previously unstudied coastal population at Fishing Bay Wildlife Management Area (MD) on the eastern shore of the Chesapeake Bay, representing the most southern breeding location sampled to date (Table S1). We also took a complete set of high resolution photographs for all birds banded in 2012-2014, using a Canon DSLR camera against a standardized color reference target (Xrite Digital SG Colorchecker), and collected standard morphological data for all banded birds.

**Table 2.**
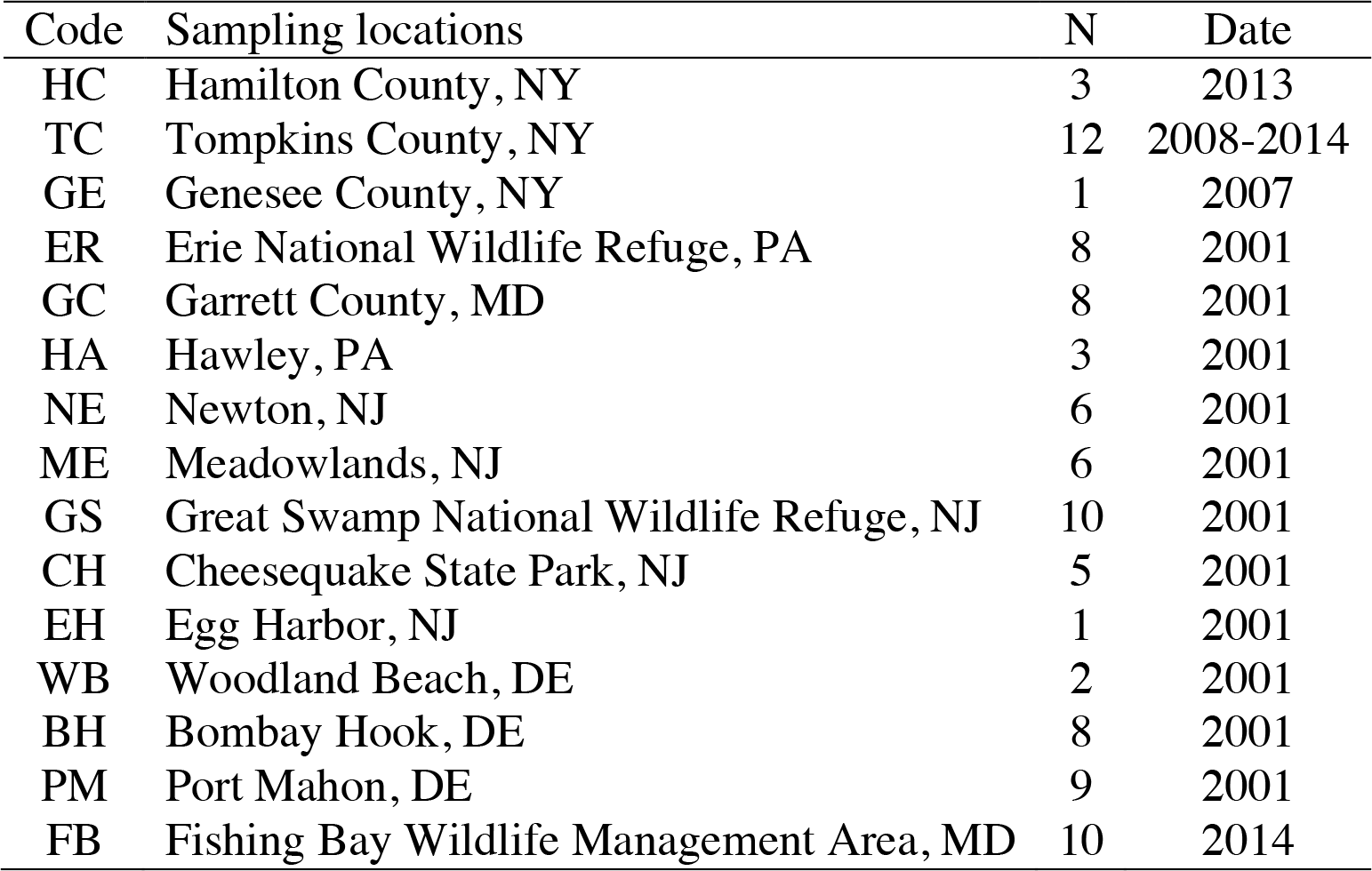
Locations and dates for Swamp Sparrows sampled in this study (N=92).

| Code | Sampling locations | N | Date |
| --- | --- | --- | --- |
| HC | Hamilton County, NY | 3 | 2013 |
| TC | Tompkins County, NY | 12 | 2008-2014 |
| GE | Genesee County, NY | 1 | 2007 |
| ER | Erie National Wildlife Refuge, PA | 8 | 2001 |
| GC | Garrett County, MD | 8 | 2001 |
| HA | Hawley, PA | 3 | 2001 |
| NE | Newton, NJ | 6 | 2001 |
| ME | Meadowlands, NJ | 6 | 2001 |
| GS | Great Swamp National Wildlife Refuge, NJ | 10 | 2001 |
| CH | Cheesequake State Park, NJ | 5 | 2001 |
| EH | Egg Harbor, NJ | 1 | 2001 |
| WB | Woodland Beach, DE | 2 | 2001 |
| BH | Bombay Hook, DE | 8 | 2001 |
| PM | Port Mahon, DE | 9 | 2001 |
| FB | Fishing Bay Wildlife Management Area, MD | 10 | 2014 |

### Library preparation and sequencing

We extracted DNA using the DNeasy kit (Qiagen), quantified concentrations on a Qubit (Invitrogen, Carlsbad, CA, USA), and balanced concentrations across samples using either dilution or vacuum filtration. We digested 175ng of each sample with enzymes SBf1 and Msp1. We followed standard ddRAD protocols for digestion, Illumina TruSeq primer ligation and clean-up as described by Peterson et al. (2012). We chose a combination of eight index groups and 12 barcodes, each 5 or 6 bases in length (Table S2). Prior to barcode ligation, we selected fragments from 400-600bp in size using the Pippen protocol (Sage Science, MA, USA). Barcode sequences were semi-randomized by systematically distributing individuals from each sampling location across index groups and barcode identities (Table S2). Within index groups, we ligated barcodes with either 14 or 16 cycles of PCR and visually inspected the products of these reactions on a gel to confirm that amplification error had not distorted the size distribution of fragments. We balanced DNA concentration to 2nM across index groups following quantification on a Bioanalyzer, and pooled index groups. We used two lanes of an Illumina HiSeq in Rapid Run mode for sequencing (yielding over 200 million 150bp reads), and we performed quality filtering and individual genotyping using a custom modification of the STACKS pipeline (Catchen et al. 2011). In brief, we executed process_radtags commands to demultiplex reads and used fastX tools for custom processing. We clipped reads with insert sizes short enough to contain adapter sequence, and trimmed all reads to 145bp. The first eight base pairs of each read did not meet quality standards due to low complexity across index sequences, and were also trimmed. We determined that trimming the sequence containing the enzyme cut site did not negatively affect the performance of the STACKS assembly by performing multiple parallel processing runs. In fact, this step improved the quality of the assembly by removing sections of low quality sequence and preserving the high quality remainder of each read. We caution, however, that this modified assembly approach may not work as efficiently in comparisons across a deeper divergence time than in our system, since successful assembly of reads from multiple individuals will rely on sequence similarity downstream of the cut site.

### SNP genotyping

We quality-filtered reads, retaining those with Illumina quality scores above 24, and the resulting 137bp fragments were assembled into a catalogue of 72,246 unique loci according to a minimum depth of five reads per stack, two locus mismatches allowed within individuals, and four locus mismatches allowed in the catalog. We dropped two individuals from the northern part of the inland range (TCm4, TCu1; both sampled far from the hybrid zone) prior to SNP discovery because they possessed unusual and divergent haplotypes at several loci, representing either sequencing error or gene flow from the unsampled boreal subspecies *M. g. ericrypta*. We grouped remaining individuals (N=90) into three populations for SNP discovery, according to whether they originated from the allopatric range of either the inland (sites HC, GE, TC, ER, GC) or coastal (WB, BH, PM, FB) populations, or from sites near the contact zone between the two (HA, NE, ME, GS, CH, EH). After extensive exploration of the impact of missing data on the accuracy of population genetic estimates from our data, we conservatively required that a given SNP be present in at least 50% of the individuals in each of the three populations. A total of 29,058 single-nucleotide sites on 4,256 stacks that met these criteria were polymorphic across allopatric inland, allopatric coastal, and sites near the contact zone. We selected the first SNP from each remaining locus to define a panel of independent SNPs for subsequent analyses of genome-wide variation and divergence (N=4,238 after additional SNP quality filtering). We performed an Analysis of Molecular Variance (AMOVA; Meirmans 2006) in the program ARLEQUIN (Excoffier and Lischer 2010) to assess the distribution of total genetic variation across different hierarchical categories of population structure. We extracted a range of different population genetic metrics and locus-specific allele frequencies from the output of STACKS, including F_ST_ (a relative measure of differentiation in allele frequencies; Wright 1965; Weir and Cockerham 1984) and Φ_ST_ (a relative measure of mutational distance between populations; Meirmans and Hedrick 2010). We also tested for F_ST_ outliers using Bayescan (Foll and Gaggiotti 2008).

### Characterizing genome-wide patterns of variation and divergence

To determine whether genome-wide SNP variation reflected evolutionary distinctness of swamp sparrows from freshwater and coastal marshes, we performed k-means clustering on the results of a Principle Coordinates Analysis (PCA) using Discriminant Analysis of Principle Components (DAPC) (Adegenet; Jombart et al. 2010), and performed bayesian assignment tests using STRUCTURE (Pritchard et al. 2000). We also performed k-means clustering manually in R, to confirm the results of the DAPC algorithm. To resolve the location and extent of admixture, we estimated the probability of assignment for all individuals in the dataset to the significant genetic clusters determined in the previous step, comparing the assignments generated by both DAPC and STRUCTURE.

### Testing morphological associations

We tested for significant associations between phenotypic scores for adaptive traits (bill depth, length, width, volume and extent of black crown plumage) and discriminant scores based on genome-wide SNP variation using Spearman Rank correlation (correlation coefficient = ϱ). For consistency, only birds sampled in 2001 and measured by R. Greenberg were used in these analyses. We performed analyses of covariance (ANCOVA) to test for significant associations between morphology and discriminant score for any traits that significantly scaled with weight. We also performed T-tests to assess the degree of morphological differentiation between distinct genetic clusters inferred from DAPC.

## Results

### Genome-wide differentiation and divergence

Out of the 29,058 SNPs in the complete dataset, only 11% (3,126 SNPs) showed significant allele frequency differences (corrected F_ST_ > 0) between coastal and inland swamp sparrows. Our panel of 4,238 independent SNPs also reflected overall similarity between coastal and inland genomes (median F_ST_=0.00, mean F_ST_=0.02), and sequence divergence was similarly low (median Φ_ST_=0.03, mean **Φ**_ST_=0.05). There was a long tail to the distributions of both F_ST_ and **Φ**_ST_ (max F_ST_=0.80; max **Φ**_ST_=0.77), representing SNPs with large allele frequency differences (F_ST_), and loci with large mutational distances between inland and coastal haplotypes (**Φ**_ST_), but Bayescan did not identify any of these as significant outliers. An AMOVA including only allopatric coastal or inland sites indicated that most of the total genetic variance is partitioned across individuals (76%) or across individuals within sites (19%), providing further indication that most variable sites within the genome are not structured between coastal and inland swamp sparrows. Higher variation within groups than between groups is commonly observed in investigations of recently diverged taxa (e.g., Campagna et al. 2015). A very low proportion of the variance (1.5%) could be attributed to structure between breeding sites within each population, indicating little to no role for isolation by distance or other geographic or demographic processes operating within coastal or inland ranges. Consistent with the presence of a long tail to the distributions of F_ST_ and **Φ**_ST_, however, a small subset of genome-wide variation (3%) was partitioned between coastal and inland habitats.

The program STRUCTURE did not resolve strong support for two distinct genetic groups (best K=1), representing genetic variation within swamp sparrows as a gradual cline in allele frequencies across many individuals of intermediate assignment when K=2 was enforced (Figure 3). In contrast, according to DAPC, total genomic variation within our samples was best represented by a single discriminant axis from a scaled PCA, and the lowest Bayesian Information Criterion (BIC) scores supported two genetic clusters corresponding to birds sampled from inland freshwater vs. coastal brackish marshes (Figure 3, S1). A conventional PCA with manually applied k-means clustering also robustly supported the genetic distinctness of swamp sparrows from coastal vs. inland habitats.

**Figure 3.**
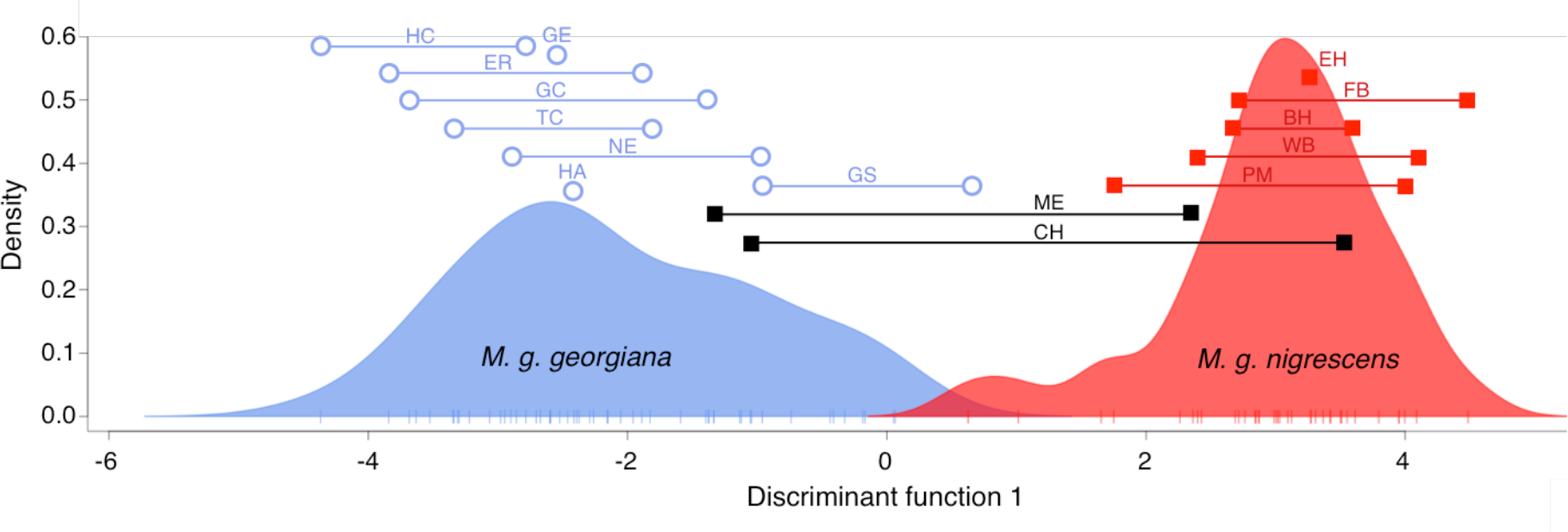
Discriminant analysis of 22 principal components, via K-means clustering and model selection, identified two clusters of genetically related swamp sparrows from a panel of 4,238 independent SNPs. A single axis of variation best distinguished the clusters. The x-axis represents the placement of individuals (short vertical lines) along this discriminant axis. The y-axis is a scaled metric representing the smoothed density of individuals corresponding to each discriminant value. Assignment probabilities consistently placed swamp sparrows from inland freshwater sites (blue) and coastal brackish sites (red) in separate clusters, with the exception of birds from two brackish sites in northern NJ (black). Individuals from these sites exhibited highly heterogeneous assignments to one cluster or the other. Brackets represent the range of genetic variation sampled from each site. Refer to Table 2 for site codes.

### Molecular variation within each population

As a group, birds sampled from coastal brackish tidal marshes possessed fewer polymorphisms across the genome than those from inland freshwater marshes (2.8% vs. 3.7% of sites), and had fewer private alleles (2,367 vs. 6,473). Average nucleotide diversity was also subtly but significantly lower in coastal birds compared to inland birds (inland π = 0.11±0.14, coastal π = 0.10±0.15; p=3.2×10^−^7), however, despite a smaller population size coastal birds exhibited more negative values of F_IS_ than inland birds (inland F_IS_=-2.0, coastal F_IS_=-3.2; p=6.3×10^−^29), suggesting that inbreeding is not responsible for this loss of variation.

### Location and extent of admixture

The two methods used to infer individual assignment probabilities (STRUCTURE and DAPC) exhibited discordance in terms of the extent of admixture, but both did exhibit concordance with respect to the location of the zone of admixture, which occurs in and around the ecotonal transition between freshwater and brackish habitats in northern NJ (Figure 4). Birds from two sites (ME and CH) had highly heterogeneous individual assignments to one cluster or the other according to DAPC, consistent with the wide range of discriminant axis values attributed to individuals from these northern brackish habitats (Figure 3, 4). Heterogeneity across individuals at these sites was also observed from the assignments calculated in STRUCTURE.

**Figure 4.**
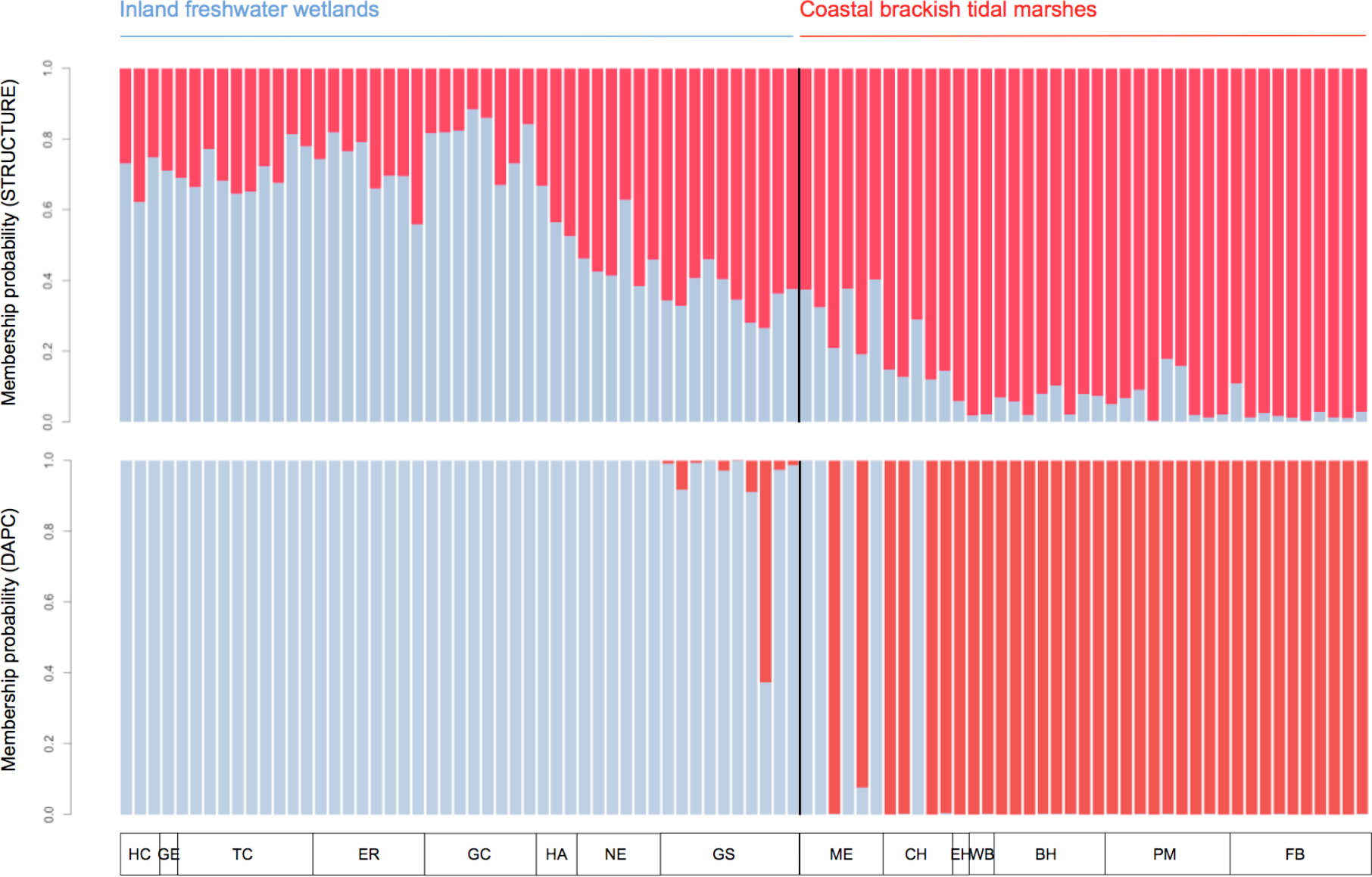
Assignment probabilities of individual swamp sparrows to one of two genetic clusters, computed by the programs STRUCTURE (top) and DAPC (bottom). Refer to Figure 1 and Table 2 for two-letter site codes. Black line indicates the ecotone between freshwater and brackish habitats.

### Associations between phenotype scores and genomic assignment

Bill length, bill width and wing chord measures were not significantly correlated with the placement of individuals on the discriminant axis (Figure S1). Correlations between DAPC discriminant axis score and morphological score were significant for the following traits: Bill depth (ϱ=0.46, p<0.001), bill volume (ϱ=0.35, p<0.05), extent of black crown plumage (ϱ=0.29, p<0.05), tarsus (ϱ=0.20, p<0.05) and weight (ϱ=0.33, p<0.05). Bill depth and volume did not significantly scale with tarsus length, a common proxy for overall size; however both metrics did scale with weight (Figure S2; bill depth, ϱ=0.41, p<0.01; bill volume, ϱ=0.49, p<0.001). We conducted analyses of covariance (ANCOVA) to test for an association between phenotype and discriminant axis score when the confounding effect of weight was removed, and both relationships remained strong and significant (p<0.001). Since variation in bill depth influences bill volume, and bill width and length were not significant, we only show the data for bill depth and black crown plumage (Figure 5). Both bill depth and black crown plumage also significantly differed between the two distinct genetic clusters inferred from DAPC (corresponding to inland and coastal sites).

**Figure 5.**
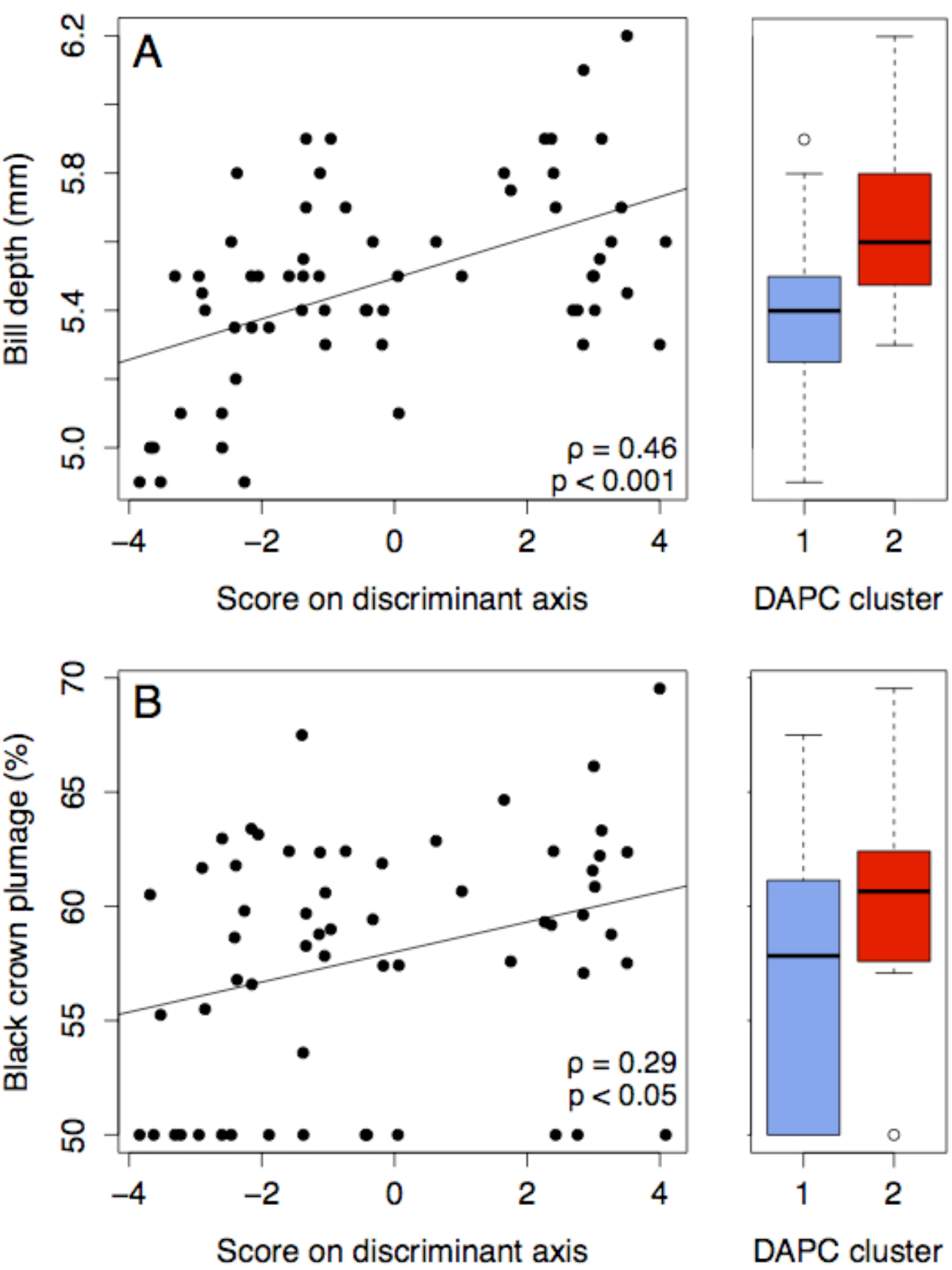
Morphological traits distinguishing inland and coastal swamp sparrows, and their association with genomic discrimination scores (left) and cluster assignment (right). Both bill depth (A) and the extent of black crown plumage (B) positively correlate with DAPC score. Birds assigned to the coastal cluster had significantly deeper bills and more extensive black crown plumage than those assigned to the inland cluster (p < 0.001 and p < 0.05, respectively).

## Discussion

### Recent adaptive divergence between inland and coastal swamp sparrows

In our comparisons, genome-wide patterns of variation supported previous evidence that inland and coastal swamp sparrows are very closely related, but have diverged very recently and are now on distinct evolutionary trajectories. We found evidence of population genetic structure corresponding to inland and coastal subspecies, but it was sufficiently low to be consistent with divergence during the most recent 15,000 years, and it was only detected by the more sensitive of the two methods of genetic cluster assignment (DAPC inferred two significant clusters but STRUCTURE did not). Across all swamp sparrows sampled, degrees of morphological variation did correlate with degrees of genome-wide differentiation, and both bill depth and black crown plumage were significantly associated with the genomic cluster to which birds were assigned.

### Divergent SNPs despite a highly similar genomic background

Inland and coastal swamp sparrow genomes are highly similar, but a long tail in the F_ST_ distribution also suggests selection in regions of the genome linked to these sites. Bayescan detected no significant outliers, however, which is not surprising given the subtle phenotypic divergence in question: Outlier tests scan for sites with reciprocally fixed or nearly fixed haplotype frequencies, and the adaptive and heritable phenotypes that differentiate inland and coastal swamp sparrows are not fixed in either population. Rather, bill depth and plumage melanism in coastal birds is an exaggerated subset of the total trait variation among inland birds (Figure 2; Greenberg and Droege 1990). Detection of functional variation responsible for adaptive phenotypes will require much more extensive genomic sampling, both to increase the proportion of the genome scanned and to enhance our ability to detect significant deviations in allele frequencies due to selection by more tightly estimating the background distribution of F_ST_ across the rest of the genome.

### Evidence of a founder effect during coastal colonization

Compared to inland swamp sparrows, coastal swamp sparrows exhibited fewer polymorphic sites, reduced nucleotide diversity at those sites, and fewer private alleles. This pattern of reduced molecular variation is not likely due to post-colonization genetic drift in the smaller coastal population since average F_IS_ is negative in coastal birds, indicating less inbreeding. In fact, coastal populations appear to exhibit even less inbreeding than the large inland population (more negative F_IS_). Reduced population genetic variation on the coast without evidence of contemporary inbreeding is therefore consistent with an alternative demographic scenario: a recent population bottleneck during coastal colonization. If coastal colonists originated from the same ancestral source population as contemporary inland populations, these founders may have carried only a subset of ancestral variation with them.

### Gene flow at the inland/coastal ecotone

Admixture estimates for individuals from across the northeastern US support earlier suggestions that there is a zone of contact between inland and coastal swamp sparrows in northern New Jersey, where inland freshwater marshes transition to brackish coastal marshes. Depending on the assignment algorithm used, this transition between genetically distinct groups is inferred to be either abrupt (DAPC) or gradual (STRUCTURE), reflecting different sensitivities to the subtle signal of divergence present in the data. Finer-scale geographic sampling of sparrows from within this region is ongoing and will facilitate the use of more computationally sophisticated methods to infer ancestry (Introgress, Gompert and Buerkle 2010; BGC, Gompert and Buerkle 2012; HapMix, Price et al. 2009; RASPberry, Wegmann et al. 2011).

### Local adaptation despite gene flow

In the present day, adaptive diversity appears to be maintained by selection despite gene flow since we detected highly admixed individuals at the ecotone between habitats in northern New Jersey. Two aspects of the swamp sparrow system may explain the maintenance of locally adaptive phenotypes despite active gene flow at that contact zone. First, since adaptive trait variation in coastal birds is a subset of the variation present inland (Figure 2), inland populations do harbor some coastally adaptive standing variation. Gene flow from the large inland population may therefore seed both adaptive and non-adaptive variants into the coastal gene pool, and if selection were strong, differential survival would mean that only the adaptive variants are maintained on the coast (eg. differential habitat performance; Harrison 1986, 1990). This would be an example of immigrant inviability maintaining adaptive diversity (Nosil et al. 2005). Second, there is robust evidence that divergent natural selection on bill size drives positive assortative mating according to habitat in swamp sparrows (Balletine 2006; Balletine et al. 2013a, 2013b). This process could maintain local adaptation by constraining gene flow to a narrow band of ecotonal or transitional habitat where divergent selection on bill shape is not strong or is not present. Future research in this hybrid zone that includes explicit tests of either mechanism will provide valuable insight into the mechanisms of adaptive evolution.

## Acknowledgements

The authors thank J. Searle, R. Fleischer and three anonymous reviewers for comments that improved this manuscript. The Smithsonian Conservation Biology Institute’s Center for Conservation and Evolutionary Genetics (R. Fleischer), and Cornell University Museum of Vertebrates (C. Dardia, V. Rohwer) contributed samples collected by N. Perlut, N. Mason and G. Seeholzer, among others. Blackwater National Wildlife Refuge (M. Whitbeck) and Fishing Bay Wildlife Management Area (J. Moulis) provided logistical support, and A. Bessler, K. Bostwick and K. Deane-Coe provided field assistance. S. Bogdanowicz and L. Campagna guided ddRAD sequencing and assembly. This research was supported by the Athena Fund at the Laboratory of Ornithology, Cornell Vertebrate Genomics, the Frank M. Chapman Fund at the American Museum of Natural History and Andrew W. Mellon Student Research fund. Fellowship support for P.D. was provided by Cornell University, the Cornell Lab of Ornithology and the Natural Sciences and Engineering Research Council of Canada (PGSD3 405451-11). Sampling was conducted under appropriate state and federal permits and all procedures conformed to protocols approved by Cornell’s Institutional Animal Care and Use Committee (IACUC). Authors Russ Greenberg and Rick Harrison passed away before the final version of this manuscript, but both were heavily involved in discussion of the ecological (R.G.) and evolutionary (R.H.) aspects of this system. They left an indelible mark on their respective fields, and are deeply missed.

